# The α-crystallin chaperones undergo a quasi-ordered co-aggregation process in response to saturating client interaction

**DOI:** 10.1101/2023.08.15.553435

**Authors:** Adam P. Miller, Susan E. O’Neill, Kirsten J. Lampi, Steve L. Reichow

## Abstract

Small heat shock proteins (sHSPs) are ATP-independent chaperones vital to cellular proteostasis, preventing protein aggregation events linked to various human diseases including cataract. The α-crystallins, αA-crystallin (αAc) and αB-crystallin (αBc), represent archetypal sHSPs that exhibit complex polydispersed oligomeric assemblies and rapid subunit exchange dynamics. Yet, our understanding of how this plasticity contributes to chaperone function remains poorly understood. This study investigates structural changes in αAc and αBc during client sequestration under varying degree of chaperone saturation. Using biochemical and biophysical analyses combined with single-particle electron microscopy (EM), we examined αAc and αBc in their apo-states and at various stages of client-induced co-aggregation, using lysozyme as a model client. Quantitative single-particle analysis unveiled a continuous spectrum of oligomeric states formed during the co-aggregation process, marked by significant client-triggered expansion and quasi-ordered elongation of the sHSP scaffold. These structural modifications culminated in an apparent amorphous collapse of chaperone-client complexes, resulting in the creation of co-aggregates capable of scattering visible light. Intriguingly, these co-aggregates maintain internal morphological features of highly elongated sHSP scaffolding with striking resemblance to polymeric α-crystallin species isolated from aged lens tissue. This mechanism appears consistent across both αAc and αBc, albeit with varying degrees of susceptibility to client-induced co-aggregation. Importantly, our findings suggest that client-induced co-aggregation follows a distinctive mechanistic and quasi-ordered trajectory, distinct from a purely amorphous process. These insights reshape our understanding of the physiological and pathophysiological co-aggregation processes of sHSPs, carrying potential implications for a pathway toward cataract formation.

## INTRODUCTION

The α-crystallins (αA- and αB-isoforms) are prototypical members of the small heat shock protein (sHSP) family of ATP-independent chaperones, with key roles in cellular proteostasis[1, 2]. sHSPs counteract detrimental protein aggregation events implicated in various human diseases, including cataract formation[3]. Humans express ten sHSP isoforms (HSPB1-10) that are differentially expressed throughout the body, functioning as holdases and serving as initial responders to diverse forms of cellular stress[4]. αA-crystallin (αAc; aka HSPB4, 19.8 kDa) and αB-crystallin (αBc; aka HSPB5, 20.2 kDa) are abundantly expressed in the eye lens, ensuring transparency for vision[5]. While αAc is primarily found in the lens, αBc exhibits ubiquitous expression, with high levels in cardiac and neuronal tissues[6]. Consequently, αBc is associated with a number of diseases, including cataract, myopathies, neuropathies, protein folding disorders (*e.g.,* Parkinson’s and Alzheimer’s), and some cancers[7, 8]. Despite their significance, our mechanistic understanding of how these sHSPs function as chaperones remains limited, due in part to their complex structural dynamics and molecular organization.

Mammalian sHSPs exhibit polydispersed oligomeric assemblies, marked by dynamic subunit interactions that are pivotal for their biological roles[9-14]. The core domain shared among sHSP subunits, the α-crystallin domain (ACD), is flanked by a variable N-terminal domain (NTD) and a flexible C-terminal domain (CTD) (**Fig. 1A**). These domains orchestrate multivalent interactions, facilitating the formation of high-order oligomers[15-23]. αAc and αBc oligomers consist of approximately 12 to 48 subunits and exhibit rapid subunit exchange dynamics[11, 12, 22, 24-28]. This dynamic behavior underpins a high degree of structural plasticity among sHSPs that is thought to enable the recognition and sequestration of a wide spectrum of destabilized client proteins. It is hypothesized that smaller assemblies or exchanging subunits are the more active states that recognize destabilized clients, while high-order oligomers are involved in sequestering and storing misfolded (or destabilized) clients[29-32]. In this stored form, the client is maintained in a refolding-competent state that may be recovered by ATP-dependent chaperones like HSP70/HSP40[33-36].

**Fig. 1.**
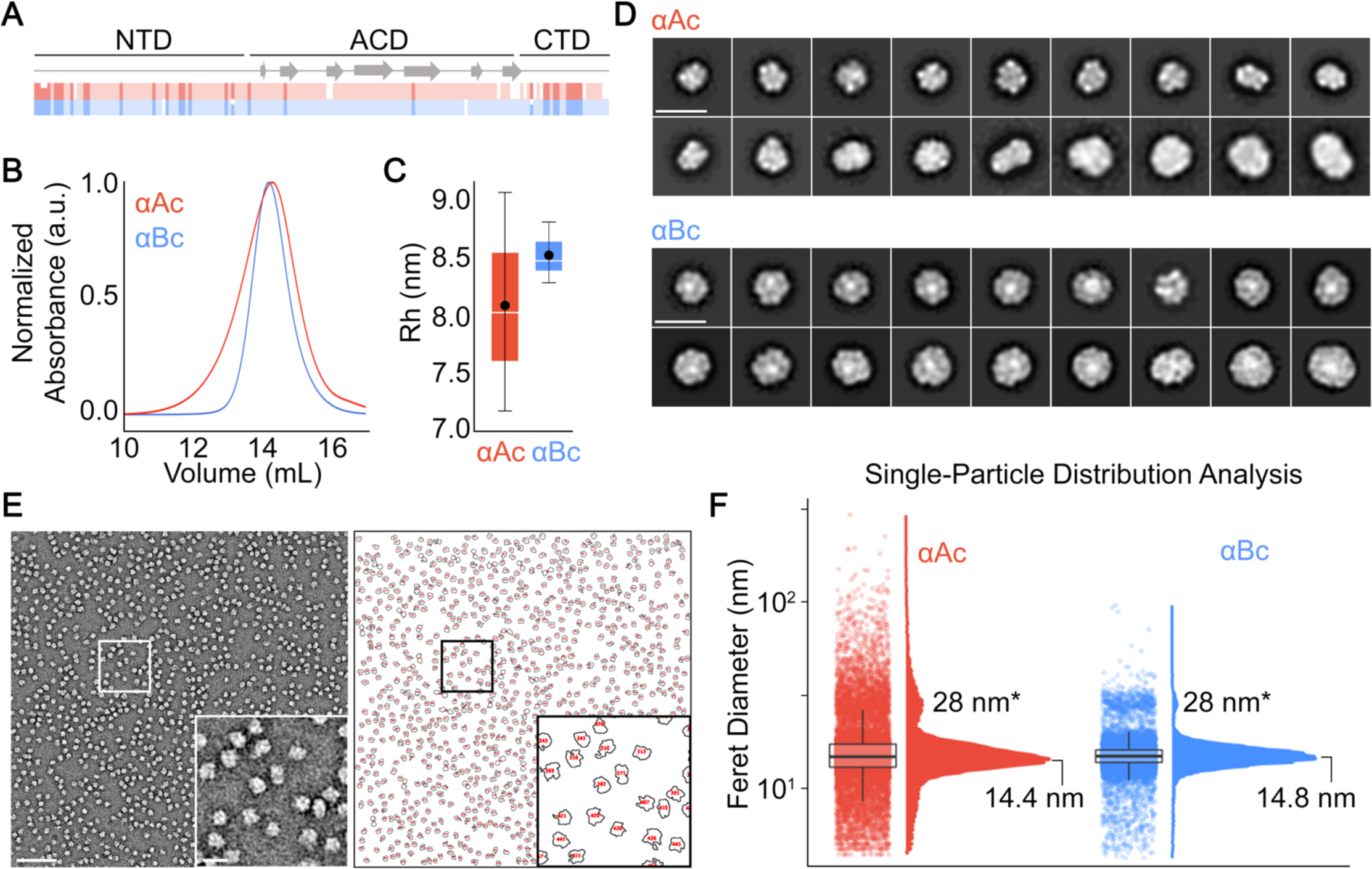
Structural analysis of polydispersed apo-states of αA- and αB-crystallin. **A.** Comparison of primary sequence homology between human αA-(αAc) and αB-crystallin (αBc), with annotations for secondary structure and domain organization. **B.** Overlay of representative size exclusion chromatography (SEC) elution profiles for αAc (red) and αBc (blue). **C.** Boxplots displaying hydrodynamic radii (R_H_) of αAc (red) and αBc (blue) determined by dynamic light scattering (n = 3). **D.** Representative 2D class averages of negatively stained αAc (top) and αBc (bottom) oligomeric assemblies (scale bar = 20 nm). **E.** Left: Representative micrograph of αBc oligomers (scale bar = 100 nm, zoom scale bar = 20 nm). Right: Representative image showing isolated particles from the same micrograph following single-particle distribution analysis using the developed FIJI workflow (see also **Supplemental Fig. 2**). **F.** Raincloud plot showing the distributions of Feret diameters of αAc (red) and αBc (blue) oligomers extracted from NS-EM images using our single-particle distribution analysis workflow shown in panel e. Feret diameters for major population modes centered around 14 nm and 28 nm are indicated.

Notably, sHSPs’ sequestration of destabilized proteins does not preclude formation of insoluble aggregates. Saturating binding conditions can lead to chaperone-client co-aggregates, while still enabling functional recovery of the client, in contrast to misfolding client aggregates formed without sHSPs[37-39]. However, the extreme degree of structural heterogeneity associated with this co-aggregation process has posed significant challenges to structural analysis and extraction of mechanistic principles associated with sHSP client sequestration and co-aggregation. In the context of the eye lens, which lacks protein turnover and ATP-dependent refolding pathways[40], the sHSP system can become overwhelmed in old age and result in formation of both soluble and insoluble light-scattering sHSP/client co-aggregates (*i.e.,* cataract), the leading cause of global vision loss[41]. While this phenomenon has been studied *in vitro*, the nature of the light-scattering aggregates has not been systematically analyzed[42-44].

Here, our objective was to quantitatively delineate the structural and morphological transformations of α-crystallin (αAc and αBc) during client-induced co-aggregation. We systematically compared αAc and αBc in their apo-states and across various stages of client-induced aggregation, employing lysozyme as a model client. To surmount the complexities arising from structural heterogeneity, we devised a single-particle electron microscopy (EM) image analysis workflow, enabling quantification of sHSP morphologies at the individual particle level. Through a combination of light-scattering methods, biochemical analysis, and direct visualization through single-particle EM, we unveiled a continuous spectrum of oligomeric states undergoing a remarkable degree of expansion and elongation of the sHSP scaffold. These changes culminated in the amorphous collapse of chaperone-client co-aggregates, which had the capacity to scatter light. The co-aggregates that are formed share striking similarity to polymeric species of α-crystallin isolated from aged lens tissue[45]. Importantly, these observations imply a mechanistic foundation characterized by a quasi-ordered co-aggregation pathway that is clearly distinct from a purely amorphous process that is shared between αAc and αBc. Our findings provide new insights into the morphological transitions of sHSPs during client-induced aggregation, shedding light on processes crucial for cellular proteostasis and age-related cataract formation.

## RESULTS

### Single-particle analysis reveals full extent of αA- and αB-crystallin polydispersity

To establish a baseline for our comparative study, we examined αAc and αBc under conditions resembling their basal apo-state. We employed size-exclusion chromatography (SEC), dynamic light scattering (DLS), and single-particle analysis using negative stain electron microscopy (NS-EM). Full-length human αAc and αBc were expressed in *E. coli* and isolated biochemically without engineered purification tags (**Supplemental Fig. 1** and **Methods**). To ensure consistency and account for environmental factors known to affect sHSP structure, we prepared purified samples in buffer at pH 7.4 supplemented with chelators (EDTA and EGTA) to remove trace amounts of divalent cations[46]. Prior to analysis, the samples were equilibrated overnight (∼16 hours) at 37° C to equilibrate exchange dynamics[12, 28]. Human αAc contains two cysteine residues (C131 and C142) and was kept under reducing conditions during purification using DTT and during functional assays using TCEP.

By size-exclusion chromatography (SEC), we observed that αAc forms slightly smaller and more polydispersed assemblies compared to αBc. The apparent molecular weights (m.w) of αAc and αBc, as determined by SEC, were approximately 440 kDa and 470 kDa, respectively. This finding is in line with the radius of hydration (R^H^) values obtained from dynamic light scattering (DLS), which measured 8.1 ± 1.0 nm for αAc and 8.5 ± 0.3 nm for αBc (avg ± sd; p = 0.5, t-test) (**Fig. 1 B-C** and **Supplemental Fig. 1**).

To further characterize their structural properties, we employed NS-EM and 2D classification methods for single-particle analysis (**Fig. 1 C-D** and **Supplemental Fig. 1**). The resulting 2D classes revealed that both αAc and αBc adopt a range of polydispersed oligomeric assemblies, exhibiting expected caged-like morphologies. Notably, αAc displayed a broader range of diameters (12.6 – 21.3 nm) within the resolved 2D classes compared to αBc (∼14.3 – 21.3 nm), although this difference was not statistically significant (**Fig. 1C**). While many of the assemblies observed in the NS-EM images appeared approximately spherical in projection, some 2D classes depicted complexes with oblique or irregular shapes. Although some of variation can be attributed to different viewing angles of the complexes; overall, these results are consistent with the ability of both αAc and αBc to adopt diverse symmetric and asymmetric oligomeric states, as described in previous EM studies[22, 23, 47].

The high degree of polydispersity and structural heterogeneity exhibited by sHSPs poses practical limitations on commonly applied single-particle 2D image classification and averaging methods. The outputs of these methods discretize continuum ensembles, which is not ideal for analyzing such systems [48-50]. To overcome these limitations, we employed a relatively simple and accessible workflow using available image processing tools in FIJI[51]. This approach enabled the extraction of structural descriptors from individual complexes (*e.g.,* diameter, circumference, surface area, etc.), without employing signal averaging methods, and allowed for quantitatively describing the polydispersity of sHSPs at the individual particle level (see **Fig. 1E**, **Supplemental Fig. 2** and **Methods**). While various structural descriptors could be obtained using this method, we focused on utilizing Feret diameter (*i.e.,* maximal particle diameter) as the primary descriptor of particle morphology in this study.

Feret diameters of individual particles were extracted from NS-EM micrographs of both αAc and αBc (**Fig. 1F**). To statistically compare the distribution of particle diameters, we employed a two-sample Kolmogorov-Smirnov test (K-S test). The population distributions of both αAc and αBc were found to be bimodal. The most populated mode for αAc was centered at 14.4 nm, slightly smaller than the primary mode of 14.8 nm observed for αBc. Both datasets exhibited a second mode centered at approximately 28 nm. While the population of αBc particles appeared as Boltzmann-like distributions, those of αAc displayed noticeable tails weighted toward smaller species and a prevalence of larger species compared to αBc. This dispersion is indicative of a greater degree of polydispersity in αAc. Consequently, the mean and standard deviation of particle diameters for αAc (17.4 ± 11.7 nm; avg ± sd) were larger than those for αBc (15.3 ± 3.5 nm), with a significant difference (p < 0.0001, K-S test). Importantly, the single-particle measurements and the extent of variation (*i.e.,* polydispersity) observed in αAc and αBc were consistent with our 2D classification results and DLS measurements obtained under solution-state conditions (**Fig. 1B**).

This single-particle distribution analysis offers a valuable advantage by providing quantitative information about rare states that were not well captured by 2D-class averaging or other ensemble measurements (SEC, DLS). This analysis revealed the presence of both very small and very large assemblies sampled by both αAc and αBc isoforms, with diameters ranging from approximately 5 nm to over 90 nm for αBc and over 250 nm for αAc (**Fig. 1F**). As noted, both αAc and αBc exhibited an additional minor, yet significant, population with Feret diameters of approximately 28 nm, which is roughly twice the diameter of the most populated modes. Upon closer examination of the raw micrographs, it became apparent that this population resulted from two oligomers positioned in close contact, which were unresolved by our segmentation approach. This observation suggests the possibility of a true population of "kissing oligomers" or it could be attributed to particle crowding on the EM grid (**Fig. 1F**, *asterisk*, see **Discussion**).

Based on these results, we concluded that our approach to single-particle distribution analysis of NS-EM images was effective at quantitatively describing and discriminating nuanced differences in sHSP structure and degree of polydispersity. With this development, we aimed to apply this constellation of methods to investigate the structural changes and dynamics of αAc and αBc in response to client interactions.

### αA- and αB-crystallin display different degrees of susceptibility to client-induced co-aggregation

For comparative functional analysis, αAc and αBc were assessed using aggregation suppression assays, monitored by light-scattering (turbidity) at 360 nm. Following the functional assays, the chaperone-client complexes were then directly subjected to structural characterization by SEC and DLS, without prior filtering or centrifugation (**Fig. 2**). We utilized lysozyme (lyso) as a model client due to its molecular weight (14.4 kDa), which is similar to the clients found in the eye lens, such as β/γ-crystallins. Unlike native clients, lysozyme readily unfolds at physiological pH (7.4) and temperature (37° C) in the presence of a reducing agent (TCEP), allowing for a controlled analysis of the structural effects induced on the chaperone complexes. This allowed us to determine the general mechanism of action of the chaperone without the complication of altering environmental conditions (such as heat or pH) that are known to affect sHSP structure[46]. It should be noted that the ratios of chaperone to client used in these experiments do not directly correlate with the stoichiometry of the formed chaperone/client complexes, as the aggregation suppression assays are conducted under non-equilibrium conditions. However, the observed dose-dependent response of chaperone concentrations suggests that at lower chaperone-client ratios, the chaperone is relatively more saturated by the binding of unfolding (or destabilized) client.

**Fig. 2.**
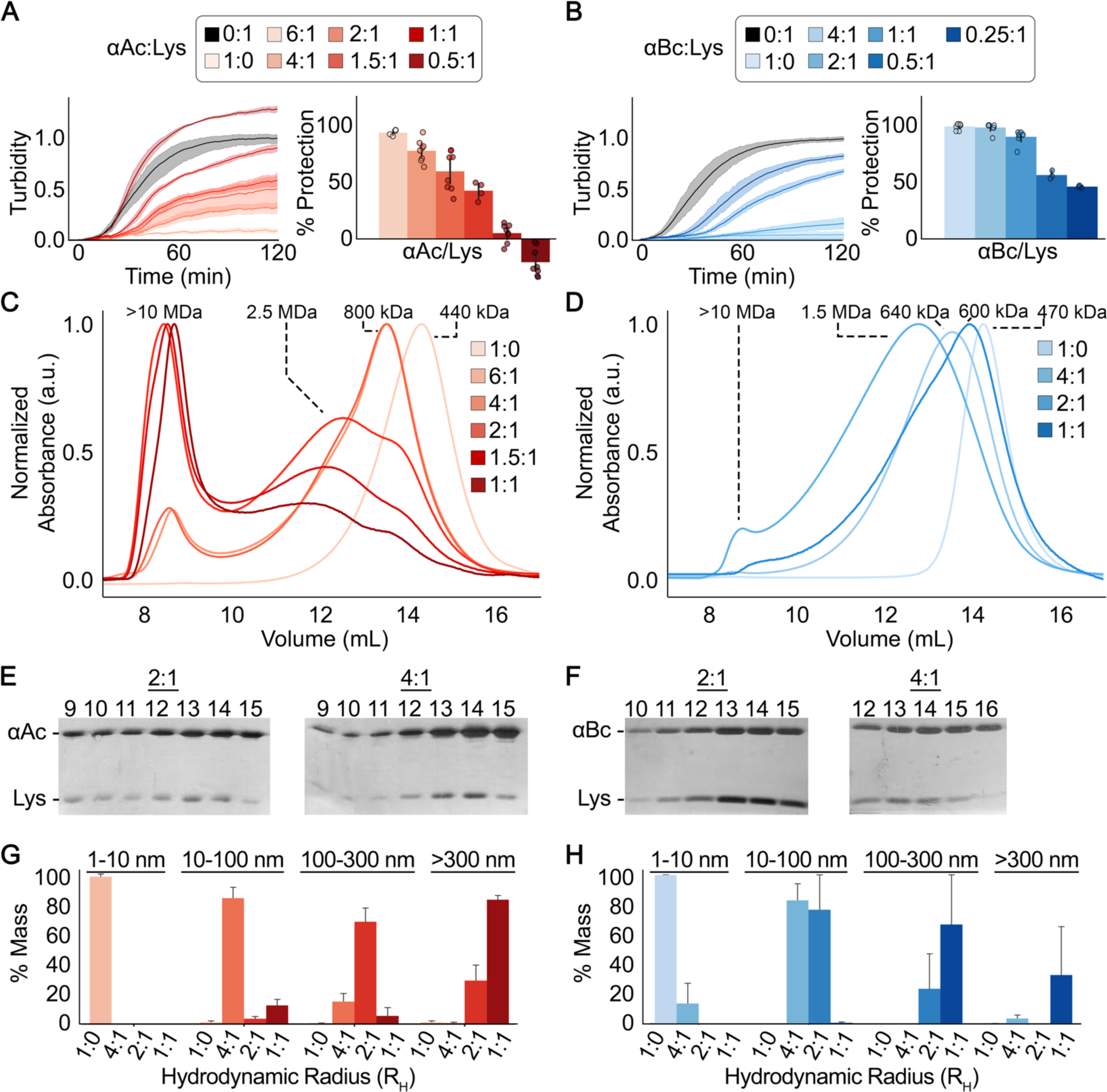
Chaperone activity and structural characterization of α-crystallin/lysozyme complexes and co-aggregates. **A,B.** Aggregation suppression assays against unfolding lysozyme (Lys) client conducted with varying molar ratios of αAc (red traces, panel A) and αBc (blue traces, panel B). Turbidity measurements at 360 nm were used to monitor aggregation. Lysozyme-only conditions (0:1 ratio) are shown in black. *Left:* Raw turbidity traces with mean +/- std displayed. *Right:* Histograms showing overall percent protection with error bars representing +/- std (n = 4 – 7). **C,D.** Overlay of size exclusion chromatography (SEC) elution profiles for αAc (red, panel C) and αBc (blue, panel D) chaperone/client complexes formed after aggregation suppression assays. The apparent molecular weights (m.w.) of major elution peaks are indicated. **E,F.** SDS-PAGE analysis of elution fractions from SEC for the 2:1 and 4:1 ratios of αAc (panel e) and αBc (panel f). Protein bands corresponding to the presence of αAc or αBc chaperone and lysozyme (Lys) are marked. **G,H.** Size distribution analysis from dynamic light scattering (DLS) measurements of chaperone/client co-aggregates formed between αAc/lysozyme (panel G) and αBc/lysozyme (panel H) at varying chaperone:client ratios of 4:1, 2:1, 1:1, and 1:0 (negative control). The measurements were taken after 2 hours of initiating aggregation suppression assays, as shown in Fig. 2A**,B**. Particle size measurements were binned at 1-10 nm, 10-100 nm, 100-300 nm, and >300 nm. Bar plots show mean +/- sem (n = 3).

Under control conditions with 10 μM lysozyme alone (0:1 ratio of chaperone:client), we observed consistent aggregation kinetics of the client, with a t_1/2_ of 33 ± 2 minutes for the αAc assay and 35 ± 3 minutes for the αBc assay (**Fig. 2A, B**; grey lines). In contrast, when αAc and αBc (pre-equilibrated at 37° C for ∼16 hours) were prepared alone and in the presence of TCEP (*i.e.,* apo-state conditions, or 1:0 ratio), no aggregation was observed under the same conditions (**Fig. 2A, B**). Furthermore, incubating αAc or αBc with lysozyme but without reducing agent also did not result in aggregation under these conditions (data not shown).

When lysozyme was treated with a reducing agent in the presence of pre-equilibrated αAc, nearly complete aggregation suppression (∼91% protection) was achieved at a molar ratio of 6:1 (chaperone:client), with a decreasing degree of protection at lower ratios (**Fig. 2A**). At a ratio of 1:1, αAc showed only ∼5% protection, but there was still a significant delay in the aggregation kinetics compared to lysozyme alone (t_1/2_ = 48 ± 3 min, p < 0.0001). Notably, at even more saturating levels of the client (0.5:1), turbidity was enhanced beyond the lysozyme-only conditions, indicated by negative protection values (-20% and **Table 1**). This result is interpreted to reflect the increased scattering caused by the formation of large co-aggregates between αAc and lysozyme (see below). In comparison, αBc exhibited higher chaperone activity against lysozyme under similar conditions, with 99% protection achieved at a ratio of 4:1 (chaperone:client) and 96% protection at a 2:1 ratio (**Fig. 2B**). At a ratio of 1:1 (where αAc showed 5% protection), αBc displayed 87% protection and still exhibited ∼17% protection even at a sub-stoichiometric ratio of 0.25:1, along with a significant delay in aggregation kinetics compared to lysozyme only conditions (t_1/2_ = 40 ± 1 min, p < 0.0001).

**Table 1.** Summary of chaperone aggregation suppression assay results. Chaperone activity of αAc and αBc toward the aggregation of reduced lysozyme (10 µM) at 37° C at various chaperone:client ratios, showing percent protection as a measure of the chaperone’s ability to suppress turbidity, number of technical replicates (n) for each ratio, and the time (minutes) indicating half aggregation (t_1/2_). Percent protection was calculated as the average ± sem turbidity of each ratio compared to the average turbidity obtained for lysozyme-only conditions for each biological replicate. Values for t_1/2_ were determined as the average ± sem time at which turbidity reached half the maximal value for each technical replicate.

| ratio | αAc/lysozyme | | αBc-lysozyme | |
| --- | --- | --- | --- | --- |
| | % protection (n) | $t_{1/2}$ (min) | % protection (n) | $t_{1/2}$ (min) |
| lysozyme | -- | $32.6 \pm 2.0$ | -- | $35.2 \pm 2.9$ |
| 6:1 | $91.4 \pm 1.3$ (4) | $29.3 \pm 3.5$ | -- | -- |
| 4:1 | $76.0 \pm 3.3$ (8) | $46.0 \pm 2.6$ | $98.8 \pm 0.2$ (7) | -- |
| 2:1 | $58.3 \pm 5.9$ (8) | $53.8 \pm 2.4$ | $95.9 \pm 1.1$ (7) | -- |
| 1.5 | $41.7 \pm 3.8$ (4) | $47.8 \pm 2.2$ | -- | -- |
| 1:1 | $5.3 \pm 2.8$ (8) | $47.6 \pm 2.9$ | $87.1 \pm 2.4$ (7) | $64.7 \pm 2.1$ |
| 0.5:1 | $-19.5 \pm 4.3$ (8) | $36.7 \pm 3.2$ | $31.9 \pm 2.0$ (7) | $64.8 \pm 2.4$ |
| 0.25:1 | -- | -- | $17.4 \pm 3.3$ (7) | $60.0 \pm 0.9$ |

Representative conditions from the completed chaperone assays were then subjected to SEC to demonstrate binding and characterize changes in structure and/or polydispersity at different chaperone-client ratios. When αAc was present at higher chaperone ratios (6:1 and 4:1), the major SEC peak shifted to a higher apparent molecular weight (∼800 kDa) compared to the apo-state (∼430 kDa). Additionally, a small left-hand shoulder peak appeared at approximately 2.5 MDa, and a minor void peak (m.w. >10 MDa) was observed (**Fig. 2C**). At lower chaperone ratios, there was a progressive increase in peak broadening, indicating higher polydispersity, accompanied by the loss of the ∼430 kDa peak and a shift towards the ∼2.5 MDa and void peaks (**Fig. 2C**).

Under conditions where αBc provided complete protection against reduced lysozyme (4:1 ratio), the SEC profile exhibited a relatively Gaussian shape, but with a shift towards higher molecular weight (∼640 kDa compared to ∼470 kDa for the apo-state). There was also an overall increase in peak broadening, indicating increased polydispersity (**Fig. 2D**). No significant void peak was observed under these conditions, which is consistent with the complete suppression of light-scattering at 360 nm. At the lower ratio of 2:1, the major SEC peak shifted further to an average m.w. ∼1.5 MDa, accompanied by even greater peak broadening. Additionally, a small void peak was observed, consistent with minor contribution to scattering at 360 nm under these conditions.

When αBc was applied to SEC at the lowest ratio (1:1), a major peak centered at an apparent m.w. ∼600 kDa was observed, accompanied by a noticeable left-hand shoulder at m.w. ∼1.7 MDa. However, no significant void peak was observed under this condition. Initially, this result seemed unexpected, as larger molecular weight species were anticipated under more saturating client conditions. Nevertheless, this consistent observation suggests that larger aggregates formed under these conditions were not entering the SEC column. This notion is supported by the lack of SEC elution profiles for samples prepared at lower chaperone:client ratios for both αAc (*e.g.,* 0.5:1) and αBc (*e.g.,* 0.5:1 or 0.25:1), as well as by the subsequent detection of very high molecular weight species by DLS and NS-EM discussed below. Analysis of the elution fractions by SDS-PAGE confirmed the presence of both lysozyme and αAc or αBc in the resolved SEC peaks (shown for the 2:1 and 4:1 ratios in **Fig. 2E,F**), indicating the formation of relatively stable and long-lived complexes under these conditions.

### Saturating client conditions induce structural transitions toward light-scattering α-crystallin/lysozyme co-aggregates

To further evaluate the structural effects of client binding, we analyzed apo-αAc, apo-αBc, and their complexes with lysozyme at representative chaperone:client ratios using DLS. The data were binned into four size ranges based on the radius of hydration (R_H_): 1-10 nm, 10-100 nm, 100-300 nm, and >300 nm (**Fig. 2G,H** and **Supplemental Fig. 3**). Under control conditions (1:0 ratios), apo-αAc and apo-αBc were predominantly within the 1-10 nm R_H_ bin (97.4 ± 1.9% and 99.8 ± 0.2%, respectively; avg ± sem). At a 4:1 chaperone:client ratio, both αAc and αBc showed the emergence of larger species and increased polydispersity, as indicated by populations in multiple R_H_ bins compared to the apo-states. The R_H_ distribution of αAc/lysozyme complexes under these conditions was 82.3 ± 8.6% (10-100 nm), 14.9 ± 8.3% (100-300 nm), and 0.7 ± 0.7% (>300 nm). Similarly, for αBc, there was a major population in the 10-100 nm bin, but also a persistence of species in the 1-10 nm bin, with populations represented by R_H_ values of 13.7 ± 13.7% (1-10 nm), 82.8 ± 13.0% (10-100 nm), and 3.6 ± 2.8% (>300 nm) under the same conditions.

At a 2:1 ratio, αAc and αBc populations are both shifted to larger species. The majority of αAc/lysozyme complexes are found in the 100-300 nm bin (67.7 ± 15.3%), with a significant population in the >300 nm bin (28.8 ± 16.9%). Under these conditions, αBc/lysozyme complexes are predominantly in the 10-100 nm bin (76.5 ± 23.5%), but with a notable population in the 100-300 nm category (23.5 ± 23.5%). At a 1:1 ratio, the majority of αAc/lysozyme complexes have an R_H_ >300 nm (82.3 ± 3.7%), consistent with significant light scattering at 380 nm. Only small populations are observed in the 10-100 nm (12.3 ± 6.0%) and 100-300 nm bins (5.5 ± 5.5%). For αBc/lysozyme complexes under these same conditions, significant populations are also found at >300 nm (32.6 ± 32.6%), but the most populated states fall into the 100-300 nm bin (66.7 ± 33.3%). The presence of >300 nm species in αBc is consistent with the light scattering observed at 360 nm in the turbidity assays. These findings support the idea that the SEC profile obtained under this condition is anomalous, likely due to the selective removal of larger species by the column filter.

### Single-particle analysis by EM reveals an expansion, elongation and amorphous collapse chaperone mechanism

We next proceeded to apply our single-particle EM analysis workflow, as described above, to obtain a more detailed and quantitative morphological characterization of the client-induced aggregation states of αAc and αBc (**Fig. 3, 4**and **Table 2**). To statistically compare the cumulative Feret diameter distributions between different chaperone:client ratios, we employed the Kolmogorov-Smirnov test (K-S test, two-sample).

**Figure 3:**
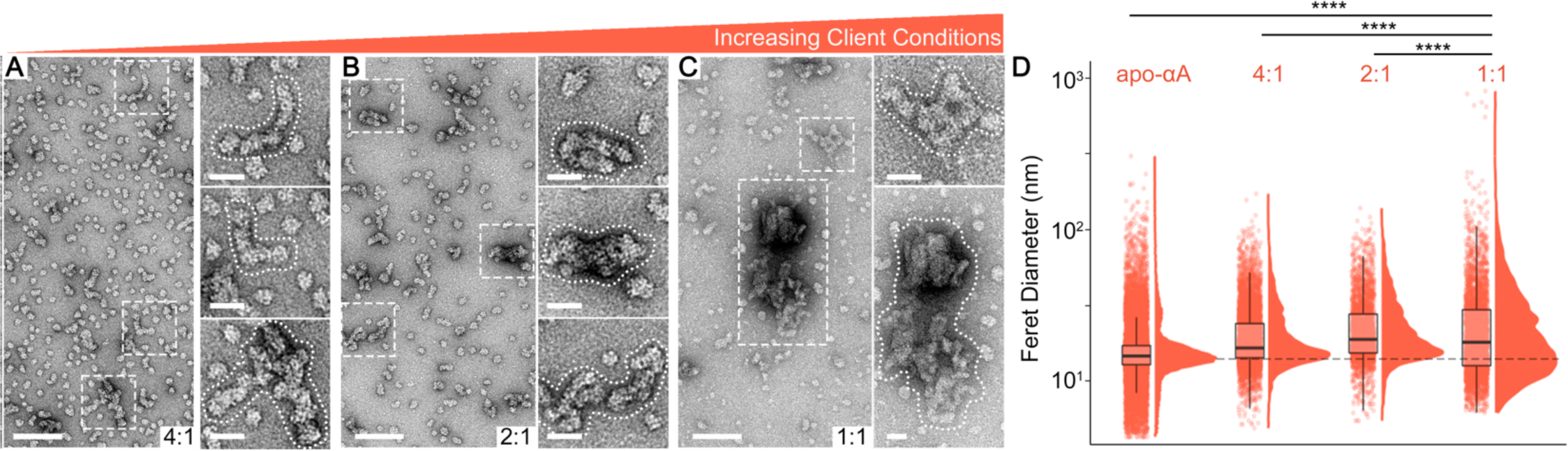
Quantitative structural analysis of αAc/Lysozyme complex formation and co-aggregation by single-particle EM. **A-C.** Representative micrographs of negatively stained αAc/lysozyme complexes and co-aggregates formed at varying chaperone:client ratios of 4:1 (panel A), 2:1 (panel B), and 1:1 (panel C), scale bar = 100 nm. The measurements were taken after 2 hours of initiating aggregation suppression assays, as shown in **Fig. 2A**. Example co-aggregates from each condition are boxed and zoomed (insets, scale bar = 25 nm). **D.** Raincloud plot showing single-particle distributions of Feret diameters extracted for apo-αAc (control) and various chaperone/client complexes and co-aggregates isolated from NS-EM datasets for the 4:1, 2:1, and 1:1 chaperone:client ratios. KS test, **** p < 0.0001

**Figure 4:**
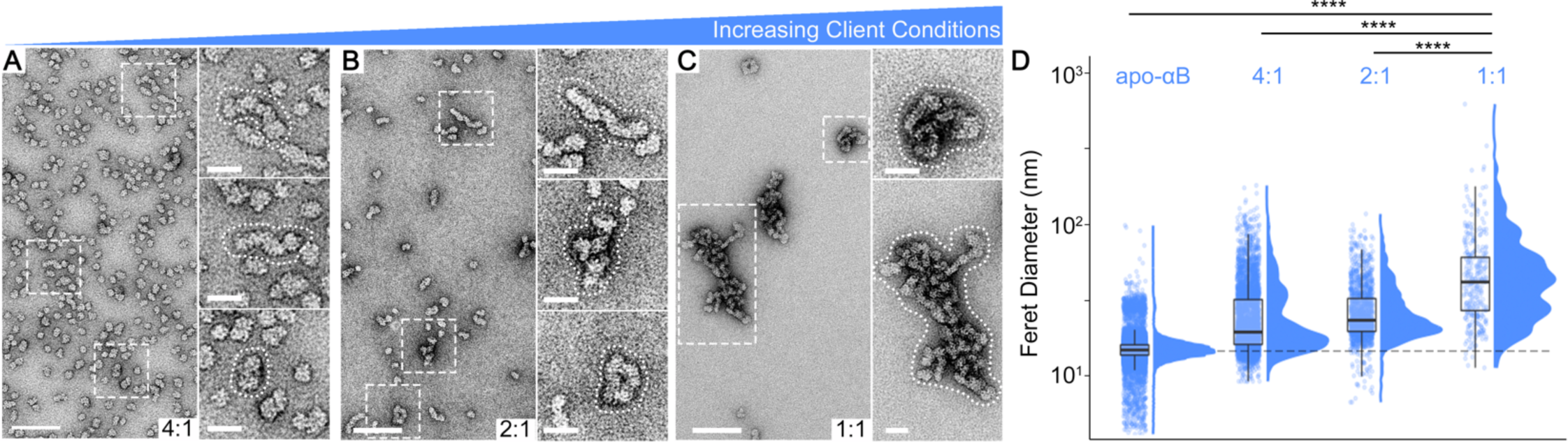
Quantitative structural analysis of αBc/Lysozyme complex formation and co-aggregation by single-particle EM. **A-C.** Representative micrographs of negatively stained αBc/lysozyme complexes and co-aggregates formed at varying chaperone:client ratios of 4:1 (panel A), 2:1 (panel B), and 1:1 (panel C), scale bar = 100 nm. The measurements were taken after 2 hours of initiating aggregation suppression assays, as shown in **Fig. 2B**. Example co-aggregates from each condition are boxed and zoomed (insets, scale bar = 25 nm). **D.** Raincloud plot showing single-particle distributions of Feret diameters extracted for apo-αAc (control) and various chaperone/client complexes and co-aggregates isolated from NS-EM datasets for the 4:1, 2:1, and 1:1 chaperone:client ratios. KS test, **** p < 0.0001

**Tabel 2.**
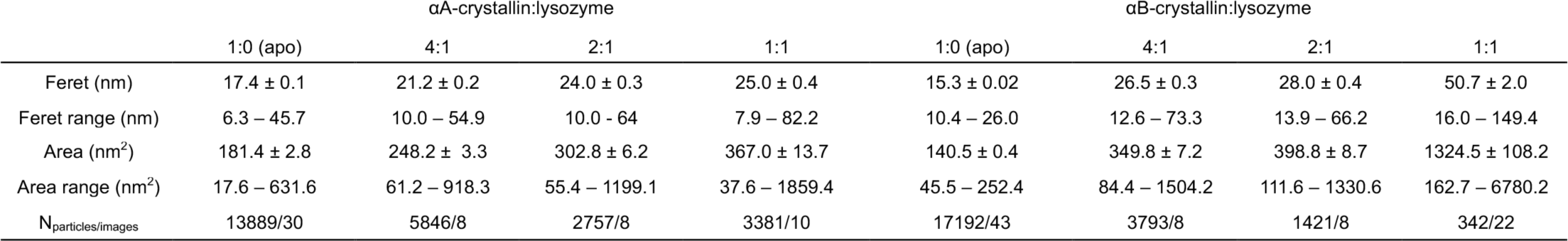
Summary of individual particle shape measurements extracted from NS-EM micrographs. Feret diameter and Area values represent the mean ± sem obtained from measurements for individual apo-state oligomers (1:0) and α-crystallin/lysozyme complexes and co-aggregates obtained at chaperone:client ratios of 4:1, 2:1 and 1:1 (see also **Fig. 3,4**) using the single-particle distribution analysis workflow described in **Methods** and **Supplemental Fig. 2**. The presented range for Feret diameter and Area distributions were calculated by truncating the smallest 2.5% and largest 2.5% of measurements to remove potential outliers.

NS-EM datasets were analyzed for αAc/lysozyme complexes formed at 4:1, 2:1, and 1:1 chaperone:client ratios (**Fig. 3**). At the higher chaperone ratio of 4:1 (representing the least saturating client conditions imaged by NS-EM), it was evident that the sequestration of lysozyme led to a variety of polymorphic structures. These structures appeared as enlarged spherical-like complexes with diameters ranging from approximately 20 to 50 nm, with average Feret diameter of 21.2 ± 0.2 nm (avg ± sem) being significantly larger than the apo-αAc structures (17.4 ± 0.1, p < 0.0001; K-S test) (**Fig. 3A,D**). In addition, an emergence of elongated structures, measuring >100 nm long, were also observed. Morphologically, these largest complexes appeared consistent with the formation of highly elongated and/or daisy-chained chaperone complexes (see **Fig 3A**, *insets*).

At the more saturating 2:1 chaperone:client ratio, the formation of expanded and elongated αAc/lysozyme complexes was even more pronounced (**Fig. 3B,D**). These complexes exhibited an average Feret diameter of 24.0 ± 0.3 nm, which was significantly larger than both the apo-state and the 4:1 condition (p < 0.0001; KS test) (**Fig. 3D**). Morphologically, the most elongated complexes reached lengths of approximately 125 nm before they appeared to undergo a collapse, resulting in the formation of larger and seemingly more amorphous aggregates (see **Fig. 3B**, *insets*).

At a chaperone-client ratio of 1:1, where αAc’s chaperone activity is significantly diminished, large amorphous aggregates become predominant, reaching sizes exceeding hundreds of nanometers and approaching 1 μm (**Fig. 3C,D**). The formation of these large complexes correlates with the significant light scattering observed at 360 nm under these conditions, surpassing the scattering observed for lysozyme alone (see **Fig. 2A**). Morphologically, these structures retain the elongated and daisy-chained features observed at higher chaperone-client ratios, indicating the collapse of highly elongated chaperone scaffolds (compare **Fig. 3B** and **3C**, *insets*). These co-aggregates exhibit distinct morphology from the plaque-like aggregates formed by lysozyme alone (refer to **Supplemental Fig. 3**). Additionally, smaller irregularly shaped species with diameters of approximately 7-12 nm are prevalent under these conditions (**Fig. 3C,D**). These smaller species resemble previously described aggregates of reduced lysozyme[52] and likely represent unbound lysozyme aggregates or also possibly unsaturated αAc/lysozyme complexes (compare to **Supplemental Fig. 3**). Notably, since αAc is no longer efficient in chaperoning at these saturated levels of client, the presence of unbound lysozyme and aggregates that evade chaperone sequestration is expected. The significant population of these smaller species contributes to a mean Feret diameter of 25.0 ± 0.4 nm in this dataset, which is comparable to the 2:1 condition. However, the increased polydispersity in this data leads to a significant difference in the distribution of Feret diameters between these two conditions (p < 0.0001; KS test).

Qualitatively, the morphological appearance of αBc/lysozyme complexes observed under different client conditions closely resembles those obtained for αAc at varying client ratios, indicating a conserved mechanism of client sequestration (**Fig. 4A**). However, quantitative comparisons reveal distinct differences that likely reflect the variations in chaperone sensitivity or potency towards the model client, lysozyme. At a 4:1 chaperone-client ratio, Feret diameter measurements demonstrate a significant increase in the polydispersity of particle size distribution compared to apo-αBc conditions (26.5 ± 0.3 nm vs. 15.3 ± 0.02 nm, respectively, p < 0.0001; K-S test) (**Fig. 4D**). Morphologically, the most prevalent structures formed under these conditions are characterized by roughly spherical oligomers similar to apo-αBc (**Fig. 4D**), with the emergence of some elongated (or daisy-chained) complexes (**Fig. 4A**, *insets*).

At a ratio of 2:1, morphological differences compared to apo-αBc become more apparent. The size and polydispersity of complexes, as indicated by distribution of Feret diameters, increases significantly compared to the 4:1 condition (28.0 ± 0.4 nm vs. 26.5 ± 0.3 nm; p < 0.0001; K-S test). In contrast to the 4:1 condition, oligomers most similar in size to apo-state appear to no longer be significantly populated (**Fig. 4D**, dotted line), consistent with our solution-state DLS data. Similar to αAc, highly elongated complexes become prevalent and are readily observed in the raw micrographs, including examples where elongated complexes appear collapsed on themselves to form more amorphous-like aggregates (**Fig. 4B**, *insets*).

Finally, when αBc was imaged under the most saturating conditions (1:1 ratio) using NS-EM, we observed the prevalence of very large aggregates with diameters reaching 100’s of nanometers (**Fig. 4C**). These aggregates also appeared as highly elongated or daisy-chained chaperones that had collapsed onto themselves, forming more amorphous-like structures. However, in contrast to αAc under the same conditions, there was a notable absence of smaller oligomers. Additionally, the size distribution of αBc/lysozyme complexes at this ratio appear multimodal with distinct peaks in the 50 – 100 nm diameter range. The comparative absence of smaller species contributed to an increased Feret diameter of 50.7 ± 37.7 nm as compared to αAc/lysozyme complexes at the same 1:1 ratio (25.0 ± 21.1 nm, p < 0.0001; K-S test). These findings are consistent with the enhanced efficiency of chaperone activity exhibited by αBc under these conditions, as indicated by the high degree of protection against light-scattering (87% protection, **Fig. 2B**).

## DISCUSSION

In this study, we demonstrated that both αAc and αBc utilize an "expansion and elongation mechanism" to accommodate increasing levels of client sequestration. This mechanism involves the formation of highly elongated chaperone-scaffolds that can reach lengths of up to ∼125 nm, followed by an ultimate collapse or folding onto themselves, resulting in the formation of larger amorphous co-aggregates reaching 1 μm in size (**Fig. 5**). This expansion and elongation process is shared by both αAc and αBc, indicating a conserved mechanism. However, we also observed isoform-specific features that correlated with the chaperone activities of these two isoforms towards the model client, lysozyme. Importantly, the quasi-ordered structural transitions observed along the client-induced co-aggregation pathway suggests that this process is mechanistically controlled and distinct from uncontrolled, purely amorphous protein aggregation. Further investigation of this mechanism could have broad implications for understanding the physiological response of sHSPs to extreme cell stress and the pathophysiology of cataract-associated light-scattering aggregate formations.

**Figure 5:**
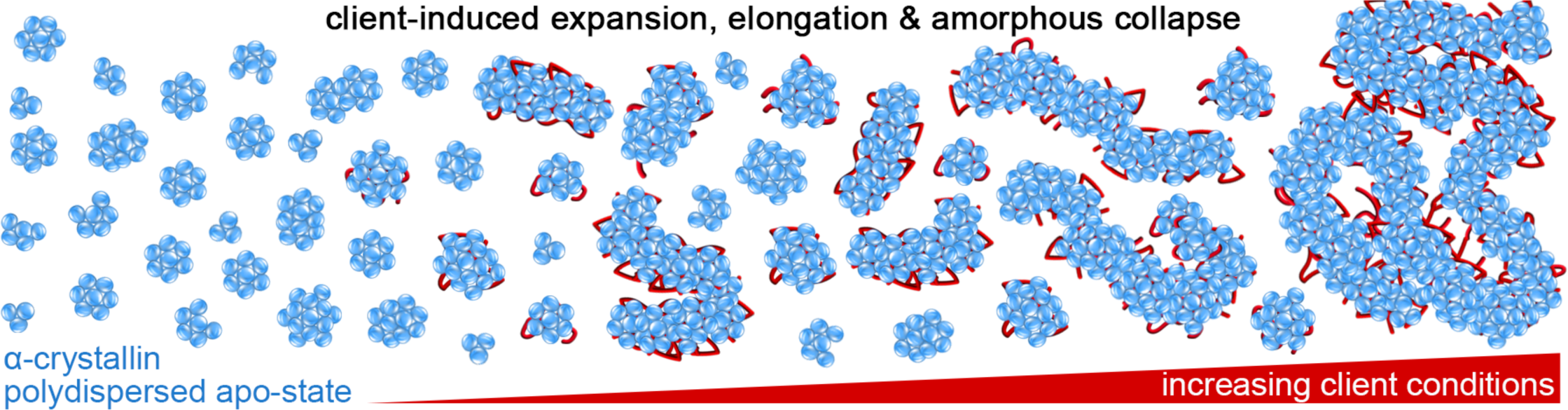
Schematic of α-crystallin client-induced expansion, elongation, and amorphous collapse. *Left:* Depiction of αAc and αBc (blue) in their apo-states represented as a polydispersed ensemble of oligomers varying in size and morphology, ranging from approximately 10 – 20 nm in diameter. *Middle to Right:* When challenged by unfolding/destabilized protein (red), αAc and αBc effectively sequester these clients, inducing structural changes seen as a complex mixture of expanded and/or elongated sHSP scaffolds. With increasing saturation of unfolding client, these chaperone/client complexes continue to elongate and extend to lengths of up to ∼125 nm before starting to collapse onto themselves, resulting in more amorphous morphologies that retain underlying features of highly elongated sHSP scaffolds. These amorphous chaperone/client co-aggregates may grow to be larger than 1 micron in size, sufficient to scatter visible light.

Under our experimental conditions, both αAc and αBc exhibited distinct morphological variability in their apo-states, with Feret diameters in the range of approximately ∼10 to ∼20 nm. Notably, αAc displayed a higher degree of polydispersity, with a wider range of smaller and larger assemblies compared to αBc. These findings align with previous structural studies by electron microscopy[9, 22, 23, 47, 53] and light scattering[54], and underscores the notion that α-crystallins can adopt a continuum of stoichiometric and quaternary structural arrangements contributing to the challenges in defining their high-resolution structures that have long eluded investigators despite decades of effort[9, 55].

The polydispersity of sHSPs is believed to be facilitated by rapid subunit or dimer exchange dynamics[12, 25-27, 55], which have led to a "traveling subunit" model to explain this mechanism. Structures corresponding to individual subunits or dimers are expected to have maximal diameters of approximately 5 – 7 nm[56-58]. While such species were not detected by SEC or DLS, our single-particle EM data did isolate a small but significant population of structures in this size range (∼4% and ∼1% for αAc and αBc, respectively). This observation further supports the strength of this single-particle approach and is consistent with the notion that isolated monomer or dimer species are not long-lived and represent only a minor population at equilibrium under the tested conditions[12]. Alternatively, or in addition, subunit or dimer exchange may occur through direct engagement between oligomers, at least under certain conditions. In this context, we also observed a significant population of apo-state αAc and αBc species with a diameter of approximately 28 nm using single-particle EM. Upon closer examination of raw micrographs, these species appeared consistent with the presence of "kissing oligomers." Although it is possible that this observation is influenced by the sample preparation conditions for NS-EM, such direct interactions between oligomers could effectively facilitate subunit exchange. Further exploration into the mechanism(s) of subunit exchange is still needed to fully resolve these possibilities.

The most notable distinction between the two α-crystallin isoforms was their differential chaperone capacity towards lysozyme, with αBc demonstrating a significantly higher efficacy compared to αAc. For instance, αBc achieved nearly complete suppression of lysozyme aggregation at a chaperone-client ratio of 2:1 (96% protection) and maintained 88% protection at a ratio of 1:1. In contrast, αAc required a ratio of 6:1 to achieve over 90% protection, and at a 1:1 ratio, the chaperone was completely overwhelmed, resulting in negative protection values. The negative values are attributed to the combination of aggregating lysozyme and the presence of large co-aggregates formed by αAc and lysozyme, a phenomenon also observed with native lens clients that had not been fully explained[42-44, 59]. Morphologically, these functional differences are reflected in αAc’s greater polydispersity and larger complex formation at lower client concentrations compared to αBc, as evidenced by SEC, light-scattering, and NS-EM analyses.

Despite differences in their chaperone capacity, both αAc and αBc exhibit similar response mechanisms when confronted with unfolded (or destabilized) client proteins. Indeed, understanding of these differences in chaperone capacity contributes to a more comprehensive understanding of the chaperone-response mechanism revealed by the structural characterizations in this study (**Fig. 5**). At conditions of high chaperone protection, such as the 4:1 chaperone-client ratio for αBc, chaperone complexes exhibit a morphology resembling the apo-state, with slightly expanded spherical-oblique caged-like structures and some elongated structures. This suggests minimal perturbation to the sHSP scaffold under minimal conditions of client sequestration. Shape analysis suggests that an overall spherical expansion of the chaperone in response to client persists up to a Feret diameter of approximately 20-30 nm, while significantly larger species take on an elongated "quasi-filament like" morphology. Under intermediate conditions of client challenge (e.g., 2:1 and 1:1 ratios for αBc or 4:1 and 2:1 ratios for αAc), highly elongated structures become predominant, and beyond an extension of approximately 125 nm, these structures collapse onto themselves, forming amorphous-like structures that can reach sizes exceeding hundreds of nanometers in diameter. Ultimately, under conditions of overwhelming client interaction, such as the 1:1 ratio condition for αAc, large chaperone-client co-aggregates of exceeding 1 μm in size are formed. In addition, NS-EM images obtained under these conditions show a prevalence of small aggregates that are expected to be un-bound lysozyme aggregates. This assertion is based on comparison to lysozyme-only controls (**Supplemental Fig. 3**) and the expectation that the chaperone’s capacity has been fully depleted under these conditions. However, we also cannot exclude the possibility of the presence of unbound (or inactive) αAc chaperone under these conditions.

The mechanism underlying the elongation of the chaperone scaffold remains unclear, as there are multiple potential pathways that could contribute to the formation of the observed structures in this study. One possibility is the formation of elongated chaperone complexes through daisy chaining, where associated chaperone complexes are connected by binding to common client proteins. Another possibility is that the chaperone undergoes oligomeric remodeling to accommodate client sequestration, involving subunit exchange. It is also conceivable that elongated chaperone complexes are formed through a combination of these mechanisms. However, it is worth noting that the elongation of the chaperone scaffold appears to have a specific directionality. Moreover, we do not observe strong evidence of branching or large-scale clustering, which would be expected if the elongation occurred randomly through a daisy-chaining mechanism alone. Notably, remarkably similar elongated morphologies for αAc and αBc have been observed upon heating[60], suggesting this structural transition is innate to the sHSP assemblies. However, a comprehensive understanding of these possibilities will require further investigation.

Overall, the findings in this study provide valuable insights into the sequestration of destabilized client proteins by the dynamic sHSP system and the formation of chaperone-client co-aggregates under conditions of overwhelming client challenge. In the context of the eye lens, the chaperone to client ratio will ultimately determine the efficiency of the sHSP system. In old age, as α-crystallins become depleted and destabilized clients become more prevalent, the balance will ultimately be pushed toward co-aggregation and formation of light-scattering opacities responsible for cataract and vision loss. Therapeutic approaches targeting the preservation of intact chaperone in the lens may therefore be beneficial in preventing this global vision problem. Moreover, the results here suggest a definable mechanistic basis for client-induced co-aggregation, which might ultimately be inhibited by pharmacological intervention. Further exploration into the universality of the expansion-elongation mechanism across different client types, sHSP systems, and stress conditions such as temperature and oxidation will undoubtedly provide a more robust understanding of the sHSP co-aggregation mechanism. The single-particle analysis methods presented in this study offer an accessible and effective approach to structurally characterize and quantitatively compare such conditions.

### Limitations of the study

While addressing the limitations of this work, it is important to note several aspects of the NS-EM analysis that could potentially impact interpretation. Specimens prepared for NS-EM are necessarily prepared under dilute concentrations, placed on a solid carbon support and undergo dehydration before imaging, which might influence the formation and morphology of the observed amorphous aggregates. Additionally, the single-particle distribution analysis workflow utilized has certain selection criteria and segmentation limitations in our experience, making it difficult to analyze extremely large aggregates (*i.e.,* greater than ∼1 μm), potentially leading to an underestimation of such species formed under highly saturated client conditions. Furthermore, this approach cannot distinguish between chaperone/client complexes and unbound client aggregates, which may be present in the most saturated conditions observed for αAc. Despite these limitations, the substantial agreement of results obtained by this approach with solution-state DLS measurements and previous studies supports its utility.

## ACKNOWLEDGEMENTS

We thank Dr. Russell McFarland for helpful discussions and the staff at the OHSU Multiscale Microscopy Core for EM instrumentation access and training. The research was funded by National Institutes of Health grants R01EY030987 and R35GM124779 (to S.L.R.), R01EY027012 (to K.J.L.), and T32EY23211 and F31EY033226 (to A.P.M.).

## AUTHOR CONTRIBUTIONS

A.P.M. and S.E.O. performed experiments; S.L.R. and K.J.L. conceived this study, with contribution from all authors; A.P.M. and S.L.R. wrote the original draft and all authors contributed to the final manuscript.

## METHODS

### Expression and purification of recombinant αA- and αB-crystallin

Expression and purification of αAc and αBc was adapted from Horwitz[1]. Recombinant human αAc and αBc in the expression vector pET3d were heterologously expressed in *E. coli* BL21(DE3) cells. Cells were grown at 37° C in LB media supplemented with ampicillin until reaching OD_600_ of 0.7 – 1.0. For αAc, expression was induced with 1 mM isopropyl β-D-1-thiogalactopyranoside (IPTG) followed by overnight expression at 18° C. For αBc, expression was induced with 1 mM IPTG for four hours at 37° C. Cells were harvested by centrifugation at 15,000 x g for 15 min at 4° C and resuspended in lysis buffer, containing: 20 mM Tris-HCl (pH 7.4 for αAc and pH 8.0 for αBc) aliquoted and frozen at -20° C.

For purification, a freshly thawed cell suspension was supplemented with 0.5 mM 1,4-dithiothreitol (DTT) and 0.1 mM PMSF, lysed by sonication, and cleared by ultracentrifugation at 165,000 r.c.f. for 30 min at 4° C to remove cellular debris. The supernatant was treated with DNase I (∼400 units, Thermo Scientific) for 30 min on ice and passed through a 0.22 µm filter. The clarified lysate was loaded onto a gel filtration column packed with sephacryl 300 resin (S-300; Pharmacia) equilibrated in 20 mM Tris-HCl and 1 mM EDTA (pH 7.4 for αAc and pH 8.0 for αBc). Gel filtration fractions were assessed by SDS-PAGE and fractions containing αAc or αBc were pooled and supplemented with 0.5 mM DTT. The pooled fractions were further purified by anion exchange chromatography (MonoQ; GE Healthcare) equilibrated in buffer containing: 20 mM Tris-HCl, 1 mM EGTA and 0.16 mM EDTA (pH 7.4 for αAc and pH 8.0 for αBc) and eluted with a 1 M NaCl gradient. Fractions pertaining to the elution peak for αAc or αBc were pooled, concentrated to 2 mL using a centrifugal device (Vivaspin, 100,000 kDa m.w.c.o.), and loaded onto a Superose 6 size exclusion chromatography (SEC) column (GE Healthcare) equilibrated in 20 mM HEPES (pH 7.4), 100 mM NaCl, and 1 mM EDTA. Fractions containing αAc or αBc were pooled and concentrated to ∼60 – 100 mM with a 100,000 m.w.c.o. spin concentrator (Vivaspin), aliquoted, flash frozen in liquid nitrogen, and stored at -80° C. Protein purity was assessed by SDS-PAGE and concentrations were determined by UV absorbance at 280 nm using the extinction coefficients 16,507 M^-1^cm^-1^ (αAc) and 19,000 M^-1^cm^-1^ (αBc). Due to the tendency of α-crystallin to co-purify with nucleic acids the ratio of A280/A260 was determined as >1.5 on purified samples. To maintain αAc in a reduced state, 0.5 mM DTT was added to purified protein prior to flash freezing and storage at -80 °C.

### Chaperone aggregation suppression assays

Aggregation assays were performed in a Nunc clear bottom 384 well plate (ThermoScientific) in 20 mM HEPES, 100 mM NaCl, 1 mM EDTA (pH 7.4) and 2mM tris(2-carboxyethyl)phosphine (TCEP). Lysozyme (Fisher, MS grade) was used as an unfolding client and held at a constant concentration of 10 µM for all reactions. Freshly purified samples of αAc and αBc were incubated at 37° C overnight to equilibrate quaternary structure prior to performing the aggregation assays. The reduction induced aggregation of 10 µM lysozyme was monitored in the absence and presence of varying chaperone concentrations between 60 – 5 µM for αAc (*i.e.,* chaperone:client ratios of 6:1, 4:1, 2:1, 1.5:1, 1:1, and 0.5:1) and 40 – 2.5 µM for αBc (*i.e.,* chaperone:client ratios of 4:1, 2:1, 1:1, 0.5:1, and 0.25:1). Chaperone/client mixtures were incubated at 37° C for 15 min followed by the addition of TCEP to initiate lysozyme unfolding. Turbidity at 360 nm was measured on a Tecan Infinite M200 Pro with a constant temperature of 37° C for 120 minutes. Aliquots of pooled aggregation reactions were set aside for SDS-PAGE and negative stain EM (NS-EM). The remainder of the pooled reactions were loaded onto a Superose 6 SEC column (GE Healthcare) equilibrated with 20 mM HEPES (pH 7.4), 100 mM NaCl, and 1 mM EDTA to assess the size of soluble chaperone/client co-aggregates. The pooled reactions were not filtered or centrifuged prior to SEC analysis. The Superose 6 column (24 mL bed volume) was calibrated using a commercial calibration kit (Biorad) containing thyroglobulin (670 kDa), g-globulin (158 kDa), Ovalbumin (44 kDa), myoglobin (17 kDa), vitamin B12 (1.35 kDa). The Superose 6 void volume was determined using dextran blue 2000. Aliquots of each size exclusion fraction were taken for SDS-PAGE to determine to co-elution profile of the chaperone and client.

Statistical analysis comparing αAc and αBc chaperone activity toward lysozyme was performed in excel and all visual interpretation of the data was done using matplotlib based libraries in python. Raw turbidity data was processed by subtracting the baseline of each replicate and normalizing to the average turbidity of reduced lysozyme replicates. Turbidity assays were replicated with 1-3 biological replicates and 4-8 technical replicates for each ratio. An F-test was performed comparing isoforms at each ratio, followed by a two-sample t-test. Aggregation half-times (t_1/2_) were determined as the time point corresponding to half maximal absorbance for each replicate. Statistical comparison of t_1/2_ values between chaperone:client ratios were done with a two-sample t-test.

### Dynamic light scattering measurements

All DLS measurements were done at 37° C in an Aurora 384 well plate on a Wyatt DynaPro plate reader III (Wyatt Technology, Santa Barbara, USA) equipped with an 830 nm laser and 150° DLS detector angle. Light scattering measurements were acquired over 10 seconds and processed in Dynamics software v7.10.1 (Wyatt). Protein samples were mixed and incubated at 37° C for 15 minutes before the addition of TCEP (2 mM final concentration) to initiate lysozyme unfolding. The aggregation of 10 µM lysozyme in the presence of αAc or αBc (40 µM, 20 µM, and 10 µM) was monitored by DLS for 120 minutes (n=3). These concentrations were the same used in turbidity analysis. Additionally, it was shown that the aggregation of 10 µM lysozyme alone was below the limit of detection on the DLS instrument and ensured that DLS measurements were monitoring the apo-state αAc and αBc oligomers and/or their co-aggregates formed with lysozyme. Due to the small size of lysozyme, control DLS measurements of reduced and oxidized lysozyme in the absence of αAc or αBc were performed at 100 µM with 20 mM TCEP to provide reliable measurement of the hydrodynamic radii within the instrument’s limit of detection.

Three technical replicates were carried out for each reaction. Hydrodynamic radii and percent mass were calculated using the Dynamics software regularization fitting algorithm for polydisperse samples. For comparative analysis, hydrodynamic radii and their corresponding percent mass were binned into one of four bins: 1-10 nm, 10-100 nm, 100-300 nm, and 300+ nm. The average and standard error of the mean were calculated for each bin of each reaction.

### Negative stain electron microscopy

For NS-EM, samples containing only purified αAc or αBc were pooled from SEC and diluted to ∼2 – 3 µM in 20 mM HEPES (pH 7.4), 100 mM NaCl, and 1 mM EDTA (αAc samples also contained 1 mM DTT). Sample grids of chaperone client mixtures at the 1:1 and 2:1 ratio were prepared directly from the aggregation reactions without dilution, while the 4:1 reactions were diluted 2x with dilution buffer. For each sample, a 3 µL drop was applied to carbon coated 400 mesh copper grids (Ted Pella) that were glow discharged at 15 mA for 1 min. Excess protein was blotted with filter paper, washed twice with ultra-pure water, stained with freshly prepared (0.75% wt vol^-1^) uranyl formate (SPI-Chem), blotted and dried with laminar air flow.

Negatively stained EM specimens were imaged on a 120 kV TEM (Tecnai T12, FEI) and micrographs were collected on a 2K x 2K CCD camera (Eagle, FEI) at a nominal magnification of 49,000 x at the specimen level. All micrographs were collected with a defocus range of ∼1.5 – 2.5 µm and calibrated pixel sizes of 4.37 Å/pixel for apo-state αBc (n = 43) and 4.401 Å per pixel for apo-state αAc (n = 30), αAc:lysozyme 4:1/2:1 (n = 19/8) and αBc:lysozyme 4:1/2:1 (n = 6/8) reactions.

### Single particle EM image analysis

Two-dimensional (2D) class averages were obtained as follows. Micrographs for apo-state αAc and αBc (equilibrated at 37° C for ∼16 hours) were pre-processed in EMAN 2.91[61] by screening for astigmatism and drift based on Thon rings of Fourier transforms following manual CTF fitting. Particles were picked with EMAN’s interactive particle picker and extracted with box sizes of 72 x 72 pixels for apo-state αAc (10405 particles) and αBc (14502 particles). The phase flipped particle stack from EMAN2.91 was normalized using relion_image_handler and imported into RELION3.0[62]. Reference-free 2D classification was performed on the CTF-corrected images in RELION3.0 using a mask size of 200 – 250 Å.

Single-particle morphological analysis developed for this work was performed in FIJI[51] using full micrographs of the apo-state αAc and αBc, 4:1, 2:1, and 1:1 reactions of αAc and αBc with lysozyme. An FFT bandpass filter was applied to each image stack using the default filter settings in FIJI (filter large structures at 40 pixels, filter small structures at 3 pixels, 5% tolerance, auto scale after filtering, saturate image when autoscaling). Next, the maximum filter was used with a default radius of 2 pixels followed by background subtraction (rolling ball radius of 25 - 50 pixels) was used on each micrograph stack. The filtered and background subtracted micrographs were subsequently binarized (with dark background). Background noise tuning of the radius parameter in the Remove Outliers tool, and erosion/dilation of binarized segments (see **Supplemental Fig. 2**).

The FIJI Analyze Particles tool was used to collect Feret diameter measurements. A search range for particle sizes was set to 90-20000 nm^2^ and particles residing on the edge of micrographs were excluded from analysis. The lower limit of 90 nm^2^ corresponds to a radius of ∼5.4 nm, which is below the range of hydrodynamic radii detected in DLS and large enough to filter out possible leftover noise following binarization. Images of numbered particle outlines were generated along with a table of particle analysis measurements. The number of particles analyzed for each sample were: 13889 for apo-αAc, 5846 for αAc:lysozyme 4:1, 2757 for αAc:lysozyme 2:1, 3381 for αA:lysozyme 1:1, 17192 for apo-αBc, 3793 for 4:1 αBc:lysozyme, and 1421 for αBc:lysozyme 2:1, and 342 for αB:lysozyme 1:1. A two-sample Kolmogorov-Smirnov test (K-S test) was performed in python (Scipy.stats) used to statistically compare the distributions of Feret diameter measurements between isoforms and at sHSP:lysozyme ratios.

### Sequence alignment of α-crystallin homologs

Amino acid sequences of αA-crystallin and αB-crystallin (human (P02489 and P02511), bovine (P02470 and P02510), and canine (P68280 and A0A8C0JXJ4)) were aligned using ClustalW in Jalview 2.11.2.7[63] and shaded based on percentage identity. Annotation of secondary structure (β-sheets) displays the consensus of three X-ray crystallographic structures of αB-crystallin (PDB 2WJ7[56], PDB 2Y1Y[57], and PDB 3L1G[58], using MODELLER[64] in ChimeraX1.15[65].

### Statistical analysis and data representation

All statistical descriptors (mean, standard deviation, standard error of the mean, mode) were calculated using the python based open-source software SciPy[66]. The presented ranges for Area and Feret diameter (Table 2) were calculated by truncating the bottom 2.5% and top 2.5% of the distribution to remove potential outliers. Two-sample Kolmogorov-Smirnov tests (K-S test) were done in SciPy. F-tests and T-tests for turbidity data were done either in Microsoft Excel or SciPy. All data plots were generated using libraries in python3, except the raincloud plots made in R Studio.

### Data Availability

Raw data used for structural analysis, electron micrographs and processed image files, are provided on zenodo doi: 10.5281/zenodo.8240041.

## SUPPLEMENTAL INFORMATION

### SUPPLEMENTAL FIGURES & LEGENDS

**Supplemental Figure 1.**
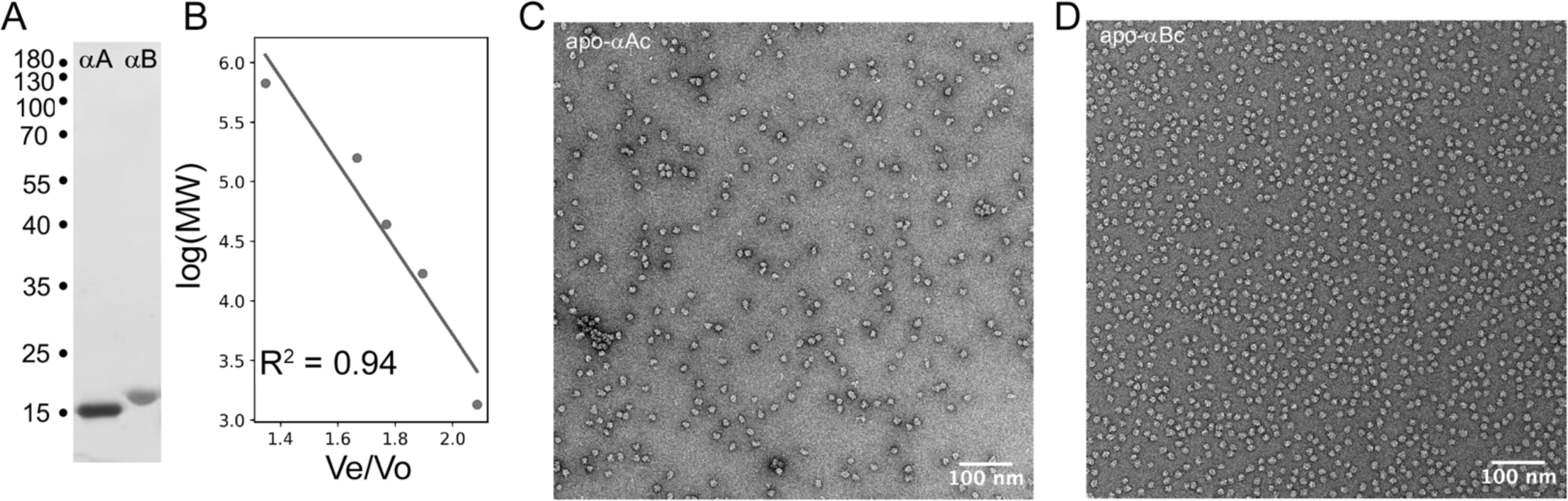
Biochemical isolation and structural assessment of recombinant apo-state α-crystallins. **A.** SDS-PAGE gel of purified αA- and aB-crystallin with molecular weight (MW) positions annotated (kDa). **B.** Standard calibration curve of Sepharose-6 size exclusion column using calibration standards (Biorad # 1511901; MW range of 1.35 – 670 kDa) and DB2000 for void volume (Vo) determination, plotted as log (MW) versus ratio of elution volume (Ve) over Vo. **C,D.** Representative NS-EM micrographs of purified apo-αAc and apo-αBc (scale bar = 100 nm).

**Supplemental Figure 2.**
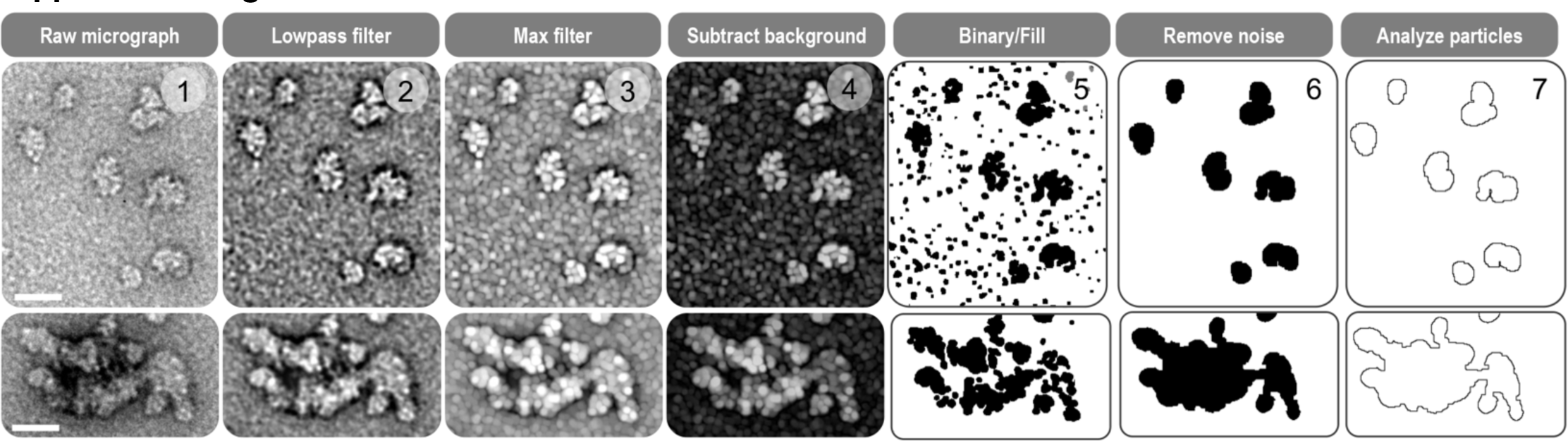
Image processing workflow applied to NS-EM micrographs for the morphological analysis of individual sHSP complexes and co-aggregates applied using the software FIJI. Example workflow shows representative co-aggregates of αAc/lysozyme at a 2:1 chaperone:client ratio. Top row of images include examples of cage-like morphologies and the bottom row shows an example of a larger co-aggregate (scale bar = 20 nm). Step 1, shows original raw micrograph with an effective pixel size of 4.40 Å/pixel. Step 2, shows result of FFT band-pass filter with applied settings: (filter large structures at 40 pixels, filter small structures at 3 pixels, 5% tolerance, auto scale after filtering, saturate image when autoscaling). Step 3, shows result of applied maximum filter set to a default radius of 2 pixels. Step 4, shows result of background subtraction (rolling ball radius of 25 - 50 pixels). Step 5, shows the result of binarization (with dark background). Step 6, shows the result of removing background noise by tuning of the radius parameter in the Remove Outliers tool, and erosion/dilation of binarized segments. Step 7, shows the final result of particle contours used for morphology analysis using the FIJI Analyze Particles tool.

**Supplemental Figure 3.**
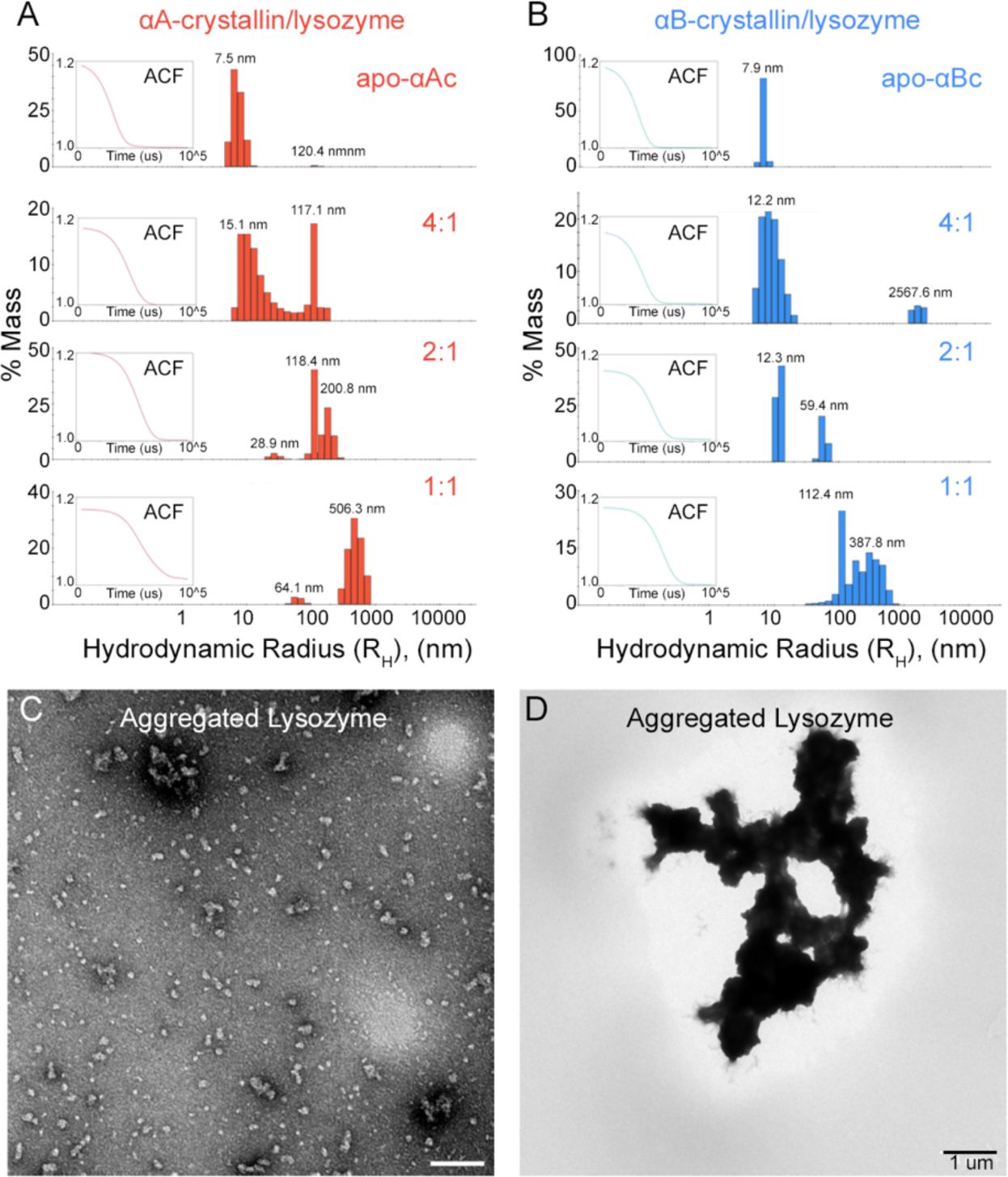
Dynamic light scattering measurements of α-crystallin/lysozyme reactions and micrograph of aggregated lysozyme. **A,B.** Mass % histogram of hydrodynamic radii (R_H_) obtained by dynamic light scattering (DLS) with autocorrelation function (ACF) for a single technical replicate shown for each experiment (inset). Representative data for αAc (panel A) and αBc (panel B) and results of aggregation suppression assays using reduced lysozyme at varying chaperone:client ratios are shown. **C,D.** Representative NS-EM micrograph of reduced lysozyme showing examples of smaller aggregates (panel C, scale bar = 100 nm) and large plaque-like aggregates (panel D, scale bar = 1 µm).

## CITATIONS

[1] Horwitz J. Alpha-crystallin can function as a molecular chaperone. Proc Natl Acad Sci U S A. 1992;89:10449–53.

[2] Wistow G. Domain structure and evolution in alpha-crystallins and small heat-shock proteins. FEBS Lett. 1985;181:1–6.

[3] Kampinga HH, Garrido C. HSPBs: small proteins with big implications in human disease. Int J Biochem Cell Biol. 2012;44:1706–10.

[4] Kappe G, Franck E, Verschuure P, Boelens WC, Leunissen JA, de Jong WW. The human genome encodes 10 alpha-crystallin-related small heat shock proteins: HspB1-10. Cell Stress Chaperones. 2003;8:53–61.

[5] Horwitz J. Alpha-crystallin. Exp Eye Res. 2003;76:145–53.

[6] Slingsby C, Wistow GJ. Functions of crystallins in and out of lens: roles in elongated and post-mitotic cells. Prog Biophys Mol Biol. 2014;115:52–67.

[7] Bakthisaran R, Tangirala R, Rao Ch M. Small heat shock proteins: Role in cellular functions and pathology. Biochim Biophys Acta. 2015;1854:291–319.

[8] Ecroyd H, Carver JA. Crystallin proteins and amyloid fibrils. Cell Mol Life Sci. 2009;66:62–81.

[9] Haslbeck M, Weinkauf S, Buchner J. Small heat shock proteins: Simplicity meets complexity. The Journal of biological chemistry. 2019;294:2121–32.

[10] Haley DA, Bova MP, Huang QL, McHaourab HS, Stewart PL. Small heat-shock protein structures reveal a continuum from symmetric to variable assemblies. J Mol Biol. 2000;298:261–72.

[11] Aquilina JA, Benesch JL, Bateman OA, Slingsby C, Robinson CV. Polydispersity of a mammalian chaperone: mass spectrometry reveals the population of oligomers in alphaB-crystallin. Proc Natl Acad Sci U S A. 2003;100:10611–6.

[12] Inoue R, Takata T, Fujii N, Ishii K, Uchiyama S, Sato N, et al. New insight into the dynamical system of alphaB-crystallin oligomers. Sci Rep. 2016;6:29208.

[13] Lambert H, Charette SJ, Bernier AF, Guimond A, Landry J. HSP27 multimerization mediated by phosphorylation-sensitive intermolecular interactions at the amino terminus. The Journal of biological chemistry. 1999;274:9378–85.

[14] Ehrnsperger M, Lilie H, Gaestel M, Buchner J. The dynamics of Hsp25 quaternary structure. Structure and function of different oligomeric species. The Journal of biological chemistry. 1999;274:14867–74.

[15] Woods CN, Ulmer LD, Guttman M, Bush MF, Klevit RE. Disordered region encodes alpha-crystallin chaperone activity toward lens client gammaD-crystallin. Proc Natl Acad Sci U S A. 2023;120:e2213765120.

[16] Klevit RE. Peeking from behind the veil of enigma: emerging insights on small heat shock protein structure and function. Cell Stress Chaperones. 2020;25:573–80.

[17] Clouser AF, Baughman HE, Basanta B, Guttman M, Nath A, Klevit RE. Interplay of disordered and ordered regions of a human small heat shock protein yields an ensemble of ’quasi-ordered’ states. eLife. 2019;8.

[18] Jehle S, Vollmar BS, Bardiaux B, Dove KK, Rajagopal P, Gonen T, et al. N-terminal domain of alphaB-crystallin provides a conformational switch for multimerization and structural heterogeneity. Proc Natl Acad Sci U S A. 2011;108:6409–14.

[19] Pasta SY, Raman B, Ramakrishna T, Rao Ch M. The IXI/V motif in the C-terminal extension of alpha-crystallins: alternative interactions and oligomeric assemblies. Mol Vis. 2004;10:655–62.

[20] Peschek J, Braun N, Franzmann TM, Georgalis Y, Haslbeck M, Weinkauf S, et al. The eye lens chaperone alpha-crystallin forms defined globular assemblies. Proc Natl Acad Sci U S A. 2009;106:13272–7.

[21] Jehle S, Rajagopal P, Bardiaux B, Markovic S, Kuhne R, Stout JR, et al. Solid-state NMR and SAXS studies provide a structural basis for the activation of alphaB-crystallin oligomers. Nat Struct Mol Biol. 2010;17:1037–42.

[22] Braun N, Zacharias M, Peschek J, Kastenmuller A, Zou J, Hanzlik M, et al. Multiple molecular architectures of the eye lens chaperone alphaB-crystallin elucidated by a triple hybrid approach. Proc Natl Acad Sci U S A. 2011;108:20491–6.

[23] Kaiser CJO, Peters C, Schmid PWN, Stavropoulou M, Zou J, Dahiya V, et al. The structure and oxidation of the eye lens chaperone alphaA-crystallin. Nat Struct Mol Biol. 2019;26:1141–50.

[24] Haley DA, Horwitz J, Stewart PL. The small heat-shock protein, alphaB-crystallin, has a variable quaternary structure. J Mol Biol. 1998;277:27–35.

[25] Bova MP, McHaourab HS, Han Y, Fung BK. Subunit exchange of small heat shock proteins. Analysis of oligomer formation of alphaA-crystallin and Hsp27 by fluorescence resonance energy transfer and site-directed truncations. The Journal of biological chemistry. 2000;275:1035–42.

[26] Aquilina JA, Benesch JL, Ding LL, Yaron O, Horwitz J, Robinson CV. Subunit exchange of polydisperse proteins: mass spectrometry reveals consequences of alphaA-crystallin truncation. The Journal of biological chemistry. 2005;280:14485–91.

[27] Delbecq SP, Rosenbaum JC, Klevit RE. A Mechanism of Subunit Recruitment in Human Small Heat Shock Protein Oligomers. Biochemistry. 2015;54:4276–84.

[28] Inoue R, Sakamaki Y, Takata T, Wood K, Morishima K, Sato N, et al. Elucidation of the mechanism of subunit exchange in alphaB crystallin oligomers. Sci Rep. 2021;11:2555.

[29] Giese KC, Vierling E. Changes in oligomerization are essential for the chaperone activity of a small heat shock protein in vivo and in vitro. The Journal of biological chemistry. 2002;277:46310–8.

[30] Shashidharamurthy R, Koteiche HA, Dong J, McHaourab HS. Mechanism of chaperone function in small heat shock proteins: dissociation of the HSP27 oligomer is required for recognition and binding of destabilized T4 lysozyme. The Journal of biological chemistry. 2005;280:5281–9.

[31] Santhanagopalan I, Degiacomi MT, Shepherd DA, Hochberg GKA, Benesch JLP, Vierling E. It takes a dimer to tango: Oligomeric small heat shock proteins dissociate to capture substrate. The Journal of biological chemistry. 2018;293:19511–21.

[32] Johnston CL, Marzano NR, Paudel BP, Wright G, Benesch JLP, van Oijen AM, et al. Single-molecule fluorescence-based approach reveals novel mechanistic insights into human small heat shock protein chaperone function. The Journal of biological chemistry. 2021;296:100161.

[33] Ehrnsperger M, Graber S, Gaestel M, Buchner J. Binding of non-native protein to Hsp25 during heat shock creates a reservoir of folding intermediates for reactivation. Embo J. 1997;16:221–9.

[34] Lee GJ, Roseman AM, Saibil HR, Vierling E. A small heat shock protein stably binds heat-denatured model substrates and can maintain a substrate in a folding-competent state. Embo J. 1997;16:659–71.

[35] Goncalves CC, Sharon I, Schmeing TM, Ramos CHI, Young JC. The chaperone HSPB1 prepares protein aggregates for resolubilization by HSP70. Sci Rep. 2021;11:17139.

[36] Wang K, Spector A. alpha-crystallin prevents irreversible protein denaturation and acts cooperatively with other heat-shock proteins to renature the stabilized partially denatured protein in an ATP-dependent manner. Eur J Biochem. 2000;267:4705–12.

[37] Haslbeck M, Miess A, Stromer T, Walter S, Buchner J. Disassembling protein aggregates in the yeast cytosol. The cooperation of Hsp26 with Ssa1 and Hsp104. The Journal of biological chemistry. 2005;280:23861–8.

[38] Cashikar AG, Duennwald M, Lindquist SL. A chaperone pathway in protein disaggregation. Hsp26 alters the nature of protein aggregates to facilitate reactivation by Hsp104. The Journal of biological chemistry. 2005;280:23869–75.

[39] Walther DM, Kasturi P, Zheng M, Pinkert S, Vecchi G, Ciryam P, et al. Widespread Proteome Remodeling and Aggregation in Aging C. elegans. Cell. 2015;161:919–32.

[40] Bloemendal H, de Jong W, Jaenicke R, Lubsen NH, Slingsby C, Tardieu A. Ageing and vision: structure, stability and function of lens crystallins. Prog Biophys Mol Biol. 2004;86:407–85.

[41] Pascolini D, Mariotti SP. Global estimates of visual impairment: 2010. Br J Ophthalmol. 2012;96:614–8.

[42] Michiel M, Duprat E, Skouri-Panet F, Lampi JA, Tardieu A, Lampi KJ, et al. Aggregation of deamidated human betaB2-crystallin and incomplete rescue by alpha-crystallin chaperone. Exp Eye Res. 2010;90:688–98.

[43] Lampi KJ, Fox CB, David LL. Changes in solvent accessibility of wild-type and deamidated betaB2-crystallin following complex formation with alphaA-crystallin. Exp Eye Res. 2012;104:48–58.

[44] Vetter CJ, Thorn DC, Wheeler SG, Mundorff CC, Halverson KA, Wales TE, et al. Cumulative deamidations of the major lens protein gammaS-crystallin increase its aggregation during unfolding and oxidation. Protein Sci. 2020;29:1945–63.

[45] Siezen RJ, Bindels JG, Hoenders HJ. The interrelationship between monomeric, oligomeric and polymeric alpha-crystallin in the calf lens nucleus. Exp Eye Res. 1979;28:551–67.

[46] Siezen RJ, Bindels JG, Hoenders HJ. The quaternary structure of bovine alpha-crystallin. Effects of variation in alkaline pH, ionic strength, temperature and calcium ion concentration. Eur J Biochem. 1980;111:435–44.

[47] Selivanova OM, Galzitskaya OV. Structural and Functional Peculiarities of alpha-Crystallin. Biology (Basel). 2020;9.

[48] Myers JB, Zaegel V, Coultrap SJ, Miller AP, Bayer KU, Reichow SL. The CaMKII holoenzyme structure in activation-competent conformations. Nat Commun. 2017;8:15742.

[49] Mostofian B, McFarland R, Estelle A, Howe J, Barbar E, Reichow SL, et al. Continuum dynamics and statistical correction of compositional heterogeneity in multivalent IDP oligomers resolved by single-particle EM. J Mol Biol. 2022;434:167520.

[50] Buonarati OR, Miller AP, Coultrap SJ, Bayer KU, Reichow SL. Conserved and divergent features of neuronal CaMKII holoenzyme structure, function, and high-order assembly. Cell Rep. 2021;37:110168.

[51] Schindelin J, Arganda-Carreras I, Frise E, Kaynig V, Longair M, Pietzsch T, et al. Fiji: an open-source platform for biological-image analysis. Nat Methods. 2012;9:676-82.

[52] Kummer N, Wu T, De France KJ, Zuber F, Ren Q, Fischer P, et al. Self-Assembly Pathways and Antimicrobial Properties of Lysozyme in Different Aggregation States. Biomacromolecules. 2021;22:4327–36.

[53] Siezen RJ, Bindels JG, Hoenders HJ. The quaternary structure of bovine alpha-crystallin. Size and charge microheterogeneity: more than 1000 different hybrids? Eur J Biochem. 1978;91:387–96.

[54] Harms MJ, Wilmarth PA, Kapfer DM, Steel EA, David LL, Bachinger HP, et al. Laser light-scattering evidence for an altered association of beta B1-crystallin deamidated in the connecting peptide. Protein Sci. 2004;13:678–86.

[55] Tardieu A. alpha-Crystallin quaternary structure and interactive properties control eye lens transparency. Int J Biol Macromol. 1998;22:211–7.

[56] Bagneris C, Bateman OA, Naylor CE, Cronin N, Boelens WC, Keep NH, et al. Crystal structures of alpha-crystallin domain dimers of alphaB-crystallin and Hsp20. J Mol Biol. 2009;392:1242–52.

[57] Clark AR, Naylor CE, Bagneris C, Keep NH, Slingsby C. Crystal structure of R120G disease mutant of human alphaB-crystallin domain dimer shows closure of a groove. J Mol Biol. 2011;408:118–34.

[58] Laganowsky A, Eisenberg D. Non-3D domain swapped crystal structure of truncated zebrafish alphaA crystallin. Protein Sci. 2010;19:1978–84.

[59] Lampi KJ, Kim YH, Bachinger HP, Boswell BA, Lindner RA, Carver JA, et al. Decreased heat stability and increased chaperone requirement of modified human betaB1-crystallins. Mol Vis. 2002;8:359–66.

[60] Burgio MR, Bennett PM, Koretz JF. Heat-induced quaternary transitions in hetero- and homo-polymers of alpha-crystallin. Mol Vis. 2001;7:228–33.

[61] Tang G, Peng L, Baldwin PR, Mann DS, Jiang W, Rees I, et al. EMAN2: an extensible image processing suite for electron microscopy. J Struct Biol. 2007;157:38–46.

[62] Zivanov J, Nakane T, Scheres SHW. A Bayesian approach to beam-induced motion correction in cryo-EM single-particle analysis. IUCrJ. 2019;6:5–17.

[63] Procter JB, Carstairs GM, Soares B, Mourao K, Ofoegbu TC, Barton D, et al. Alignment of Biological Sequences with Jalview. Methods Mol Biol. 2021;2231:203–24.

[64] Webb B, Sali A. Protein Structure Modeling with MODELLER. Methods Mol Biol. 2021;2199:239–55.

[65] Goddard TD, Huang CC, Meng EC, Pettersen EF, Couch GS, Morris JH, et al. UCSF ChimeraX: Meeting modern challenges in visualization and analysis. Protein Sci. 2018;27:14–25.

[66] Virtanen P, Gommers R, Oliphant TE, Haberland M, Reddy T, Cournapeau D, et al. SciPy 1.0: fundamental algorithms for scientific computing in Python. Nat Methods. 2020;17:261–72.

